# Speeding up the detection of invasive aquatic species using environmental DNA and nanopore sequencing

**DOI:** 10.1101/2020.06.09.142521

**Authors:** Bastian Egeter, Joana Veríssimo, Manuel Lopes-Lima, Cátia Chaves, Joana Pinto, Nicoletta Riccardi, Pedro Beja, Nuno A. Fonseca

## Abstract

Traditional detection of aquatic invasive species, via morphological identification is often time-consuming and can require a high level of taxonomic expertise, leading to delayed mitigation responses. Environmental DNA (eDNA) detection approaches of multiple species using Illumina-based sequencing technology have been used to overcome these hindrances, but sample processing is often lengthy. More recently, portable nanopore sequencing technology has become available, which has the potential to make molecular detection of invasive species more widely accessible and to substantially decrease sample turnaround times. However, nanopore-sequenced reads have a much higher error rate than those produced by Illumina platforms, which has so far hindered the adoption of this technology. We provide a detailed laboratory protocol and bioinformatic tools to increase the reliability of nanopore sequencing to detect invasive species, and we test its application using invasive bivalves. We sampled water from sites with pre-existing bivalve occurrence and abundance data, and contrasting bivalve communities, in Italy and Portugal. We extracted, amplified and sequenced eDNA with a turnaround of 3.5 days. The majority of processed reads were ≥ 99 % identical to reference sequences. There were no taxa detected other than those known to occur. The lack of detections of some species at some sites could be explained by their known low abundances. This is the first reported use of MinION to detect aquatic invasive species from eDNA samples. The approach can be easily adapted for other metabarcoding applications, such as biodiversity assessment, ecosystem health assessment and diet studies.

## Introduction

Aquatic invasive species can cause losses in biodiversity, changes in ecosystems, and impacts on economic sectors such as agriculture, fisheries, and international trade (Lovell, Stone, & Fernandez, 2006; Pimentel, Zuniga, & Morrison, 2005; Vander Zanden & Olden, 2008; Wilcove, Rothstein, Dubow, Phillips, & Losos, 1998). Traditional methods to detect aquatic invasive species, via morphological taxonomic identification of adults or larvae sampled from the environment, are often time-consuming and require a high level of taxonomic expertise. Sample processing can create a substantial delay between sample collection and completion of identifications, delaying mitigation responses (Darling & Mahon, 2011; Hatzenbuhler, Kelly, Martinson, Okum, & Pilgrim, 2017; Thomas et al., 2019)

The capture and analysis of environmental DNA (eDNA) is increasingly being conducted to detect aquatic invasive species (Ardura & Planes, 2017; Clusa, Miralles, Basanta, Escot, & García-Vázquez, 2017; Keskin, 2014; Klymus, Marshall, & Stepien, 2017; Prié et al., 2020; Rees, Maddison, Middleditch, Patmore, & Gough, 2014; Scriver, Marinich, Wilson, & Freeland, 2015; Simmons, Tucker, Chadderton, Jerde, & Mahon, 2015; Thomas et al., 2019). In general, eDNA detection approaches have been shown to overcome many of the limitations of traditional morphological approaches by decreasing the turnaround in data acquisition time, reducing costs, reducing dependency on specific taxonomic expertise and, in some cases, reducing the need to transport samples to distant laboratories by enabling species detection at, or near, the points of sampling. Most eDNA approaches to date have either employed species-specific assays using PCR, qPCR or traditional Sanger sequencing, or have targeted multiple species using Illumina-based sequencing technology. While species-specific approaches are often inexpensive and have rapid turnaround times, they are explicitly limited to a single species (or small group of species), narrowing their applicability and requiring the development of unique assays for each target species of interest, which is laborious and requires substantial validation. Multi-species approaches using Illumina-based sequencing technology overcome some of the issues of single-species approaches, but: 1) they often rely on external sequencing services, which generally have slow turnaround times of some weeks; 2) they require amassing and combining large batches of samples, to maximise sequencing efficiency; 3) sequencing cannot be performed on site and; 4) they only allow limited read lengths (c. 500 bases for Illumina-based sequencing).

More recently, a novel DNA sequencing platform, the MinION (Oxford Nanopore Technologies, UK), which uses nanopore technology, has been used to detect multiple species from eDNA samples (Truelove, Andruszkiewicz, & Block, 2019). The MinION has also been used to generate DNA barcodes and identify species from tissue-extracted DNA (Ho, Puniamoorthy, Srivathsan, & Meier, 2020; Krehenwinkel, Pomerantz, & Prost, 2019; Maestri et al., 2019; Pomerantz et al., 2018; Seah, Lim, McAloose, Prost, & Seimon, 2020). Like Illumina platforms, the MinION can be used to detect multiple species from multiple samples in a single sequencing run. Potential benefits of using the MinION include the ability to: 1) process samples without relying on external services; 2) process small batches of samples, decreasing turnaround time (because a MinION sequencing run can be stopped at any point and the flow cell reused); 3) perform on-site sequencing (the MinION is a small portable device that may be connected to a laptop computer) and; 4) sequence far longer reads (average 8 kbp for optimal sequencing). The primary potential limitation of using the MinION is that it produces higher DNA sequencing error rates. Tyler et al. (2018) recently reported an average error rate of 6 % (using R9.4 flow cells), in contrast to an average error rate of only 0.24 % observed using Illumina platforms (Pfeiffer et al., 2018), which could cause reduced reliability in species detection results. There may also be increased costs per sample and reduced scalability using MinION, although this remains to be explored in the context of species detection. If the existing limitations can be overcome, the MinION has the potential to substantially reduce sample turnaround time, thereby reducing the time-lag between obtaining samples and acquiring final results, and further increasing the usefulness of eDNA approaches for the early detection of aquatic invasive species.

In this work, we present a framework for detecting multiple invasive aquatic species using eDNA and nanopore sequencing technology. Specifically, we provide a bioinformatic pipeline that accounts for the sequencing errors produced by nanopore sequencing, thereby ensuring reliable species detection. Our approach is tested using as case study the detection of zebra mussel (*Dreissena polymorpha*) and other invasive bivalves in natural lakes and hydroelectric reservoirs. This is the first reported use of MinION to detect aquatic invasive species from eDNA samples. We show that the implementation of the framework yielded reliable species detection results in a fast, efficient and cost-effective way, thereby enhancing the value of MinION technology in molecular environmental assessment and biomonitoring.

## Materials and Methods

### Study area and target species

The field study was designed primarily to show the ability of our approach to reliably detect the highly invasive and economically-damaging zebra mussel (Pimentel et al., 2005), by comparing its detection in lentic habitats with and without previous records of the species. In addition, we also tested for the ability of the method to detect other bivalve species known to be present at each sampling site. For sites with *D. polymorpha* we selected three lakes in Italy (Lake Maggiore [212.5 km^2^]; Lake Varese [14.5 km^2^]; and Lake Lugano [48.7 km^2^]; Figure 1), where two other invasive mussels have already been detected (*Sinanodonta woodiana* and *Corbicula fluminea*). For sites without previous records of *D. polymorpha*, we selected two hydroelectric reservoirs in Portugal (Castelo de Bode [32.91 km^2^] and Caniçada [6.89 km^2^]; Figure 1). Annual monitoring of these reservoirs using conventional morphological methods (detection of larvae), have so far not detected *D. polymorpha*. Moreover, *D. polymorpha* is considered largely absent from Portuguese waters, with a single detection from a small reservoir in southern Portugal in 2019 (Catita et al., 2020).

**Figure 1.**
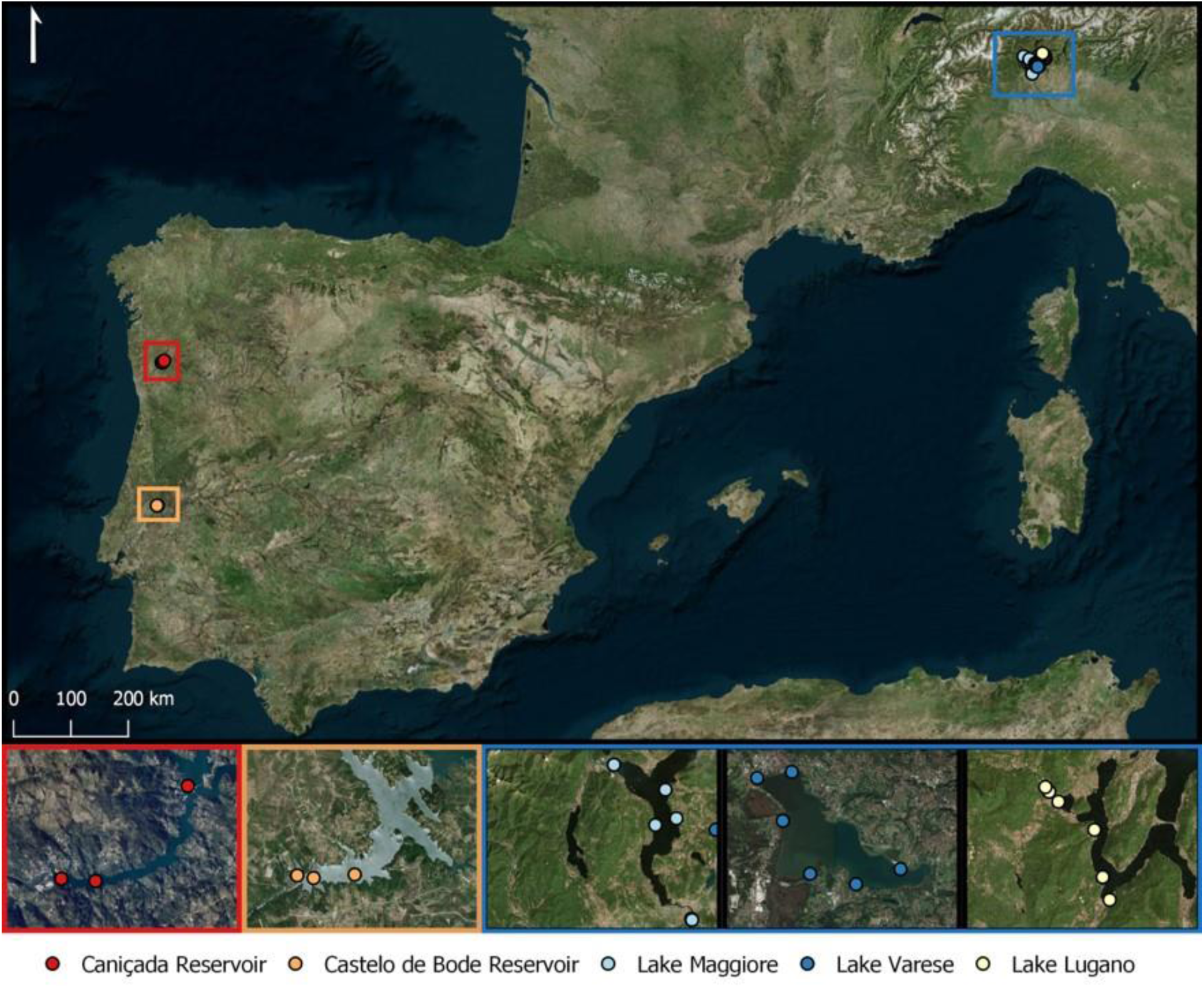
Locations of sites and sampling points used in this study, located in Portugal and Italy.

None of the species known to be present at the Italian lakes are known to be present at the Portuguese study sites, and many of the species are altogether absent from Portugal (Table 1). One species, *Margaritifera margaritifera*, was historically present at the River Cavado near the Caniçada dam site, but has not been found in recent years, although the area has been intensively surveyed (Reis, 2006; Sousa et al., 2015). This species does not occur in Italy (Lopes-Lima et al., 2017).

**Table 1.** Species expected *a priori* to be present in the sampled study areas.

| Order | Family | Species | Status in Italy | Status in Portugal |
| --- | --- | --- | --- | --- |
| Veneroida | Corbiculidae | *Corbicula fluminea*<sup>†</sup> | Invasive | Invasive<sup>‡</sup> |
| Veneroida | Dreissenidae | *Dreissena polymorpha* | Invasive | Recent invasive<sup>§</sup> |
| Unionida | Unionidae | *Sinanodonta woodiana* | Invasive | Absent |
| Unionida | Margaritiferidae | *Margaritifera margaritifera* | Absent | Native<sup>¶</sup> |
| Unionida | Unionidae | *Anodonta anatina* | Native | Native<sup>‡</sup> |
| Unionida | Unionidae | *Anodonta cygnea* | Native | Native<sup>‡</sup> |
| Unionida | Unionidae | *Anodonta exulcerata* | Native | Absent |
| Unionida | Unionidae | *Unio elongatulus* | Native | Absent |

<sup>†</sup> Note that the taxonomy of the genus *Corbicula* is controversial (e.g. Bódis, Nosek, Oertel, Tóth, & Fehér, 2011; Pigneur et al., 2011). We considered *C. largillierti* to be a lineage of *C.* *fluminea*.
<sup>‡</sup> Not known from Portuguese sites included in this study (Reis, 2006).
<sup>§</sup> A population was discovered in 2019 in southern Portugal, and is believed to have been eradicated (Catita et al., 2020).
<sup>¶</sup> Historically present at Cávado River close to the Caniçada dam, but not found in recent years (Lopes-Lima et al., 2017; Reis, 2006; Sousa et al., 2015).

### Field sampling

Field sampling was carried out during July 2019 in Italy and Portugal (Figure 1). At each lake, three to six sampling points were chosen. The choice of sampling point depended largely on accessibility to the lake. For each lake in Italy, four sampling points were chosen along the margins of the lake, as well as an additional two points located on either side of the outflow river of the lake. In Portugal, public access to the reservoir edges was very limited, so samples were taken at one point on each reservoir margin and at two points located more centrally within each reservoir, which were accessed by boat.

At each sampling point, filtration was carried out using 47 mm nitrocellulose disc filters, 0.45 µm pore size (Whatman, UK) in combination with a 500-ml filtering cup (Nalgene™ Polysulfone Filter Holder with Funnel, Thermo Scientific, USA). Water was passed through the filters using a peristaltic pump (Solinst 410, Solinst Canada Ltd., Canada) powered by a portable car battery, and silicon tubing (Solinst Canada Ltd., Canada). The target volume of water filtered was 2 L, although in some cases the filters clogged before reaching this volume. The final volume filtered for each sample was recorded. All reusable equipment (filtering cup apparatus and tubing) was sterilised between lakes by immersion in 20% bleach for at least one hour, followed by thorough rinsing. Filters were placed in 2-mL tubes with 96 % ethanol and stored at room temperature, protected from direct sunlight. Figure 2 illustrates the complete workflow of the study from sample collection to taxonomic assignment.

**Figure 2.**
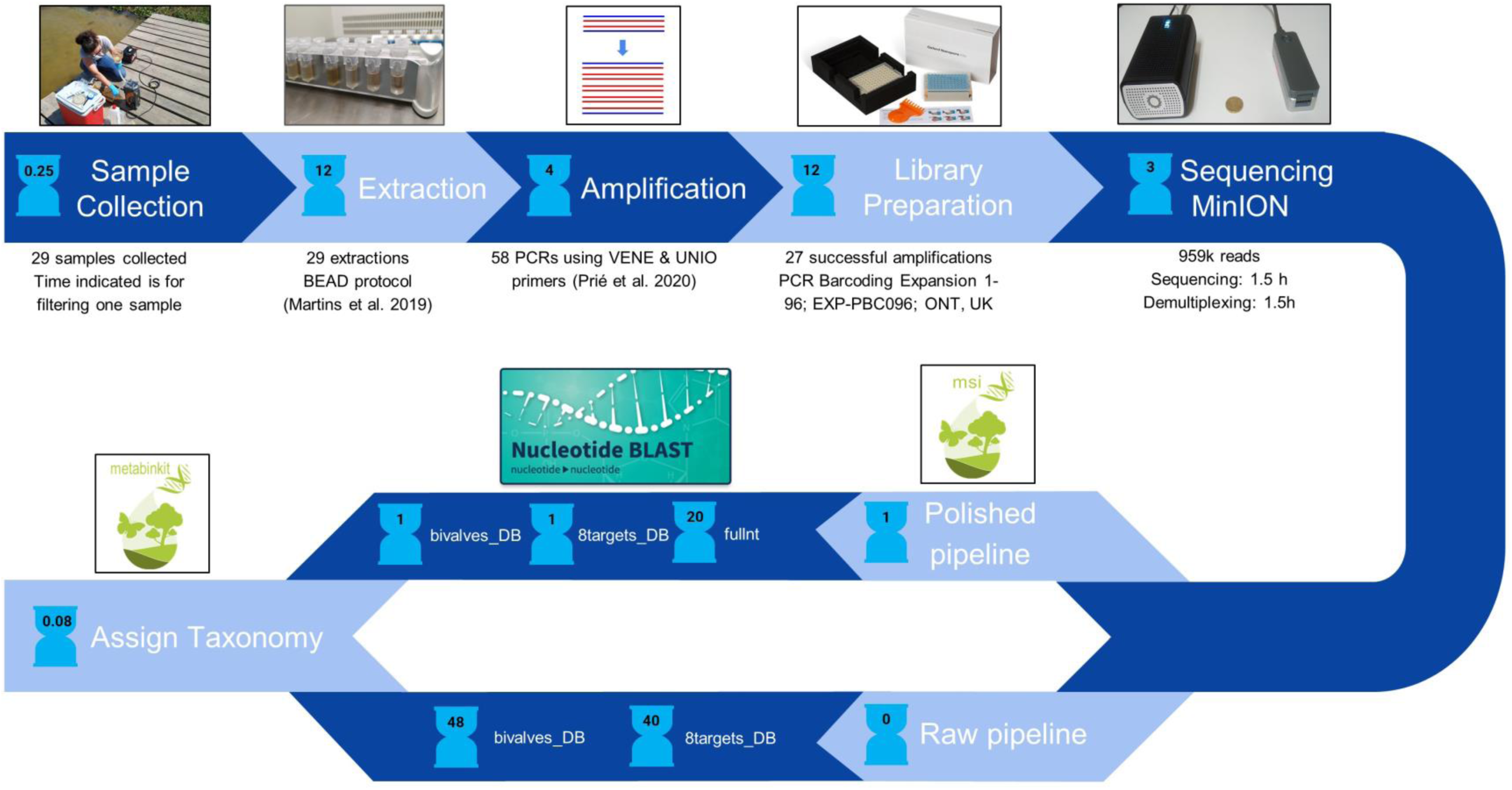
Workflow used in this study. Times indicated are in hours and include hands-on and hands-off processing times. Times indicated for bioinformatic steps are in CPU hours.

### DNA extraction

DNA extraction was performed in a positive pressure laboratory (CIBIO-InBIO, Vairão Campus, Portugal), following strict protocols which include disposable lab wear and UV sterilization of all equipment before entering the lab. DNA was extracted within eight weeks of field sampling using the BEAD protocol described by Martins et al. (2019), with the following minor modifications: the starting material consisted of half of each filter, cut into small pieces (using flame-sterilised scissors); digestion was conducted with 300 µL of ATL and 20 µL of proteinase K for 3 h at 56 °C; 100 µL of magnetic beads were used (Agencourt AMPure XP, Beckman Coulter, USA).

### PCR amplification, library preparation & sequencing

To detect eDNA originating from the order Veneroida we used the VENE primers (VENE_F 5’-CSCTGTTATCCCYRCGGTA-3’; VENE_R 5’-TTDTAAAAGACGAGAAGACCC-3’; Prié et al., 2020), and to detect species from the order Unionidae we used the UNIO primers (UNIO_F 5’-GCTGTTATCCCCGGGGTAR-3’; UNIO_R 5’-AAGACGAAAAGACCCCGC-3’; Prié et al., 2020). Primers were ordered (Eurofins Genomics, Germany), with 5’ adaptor sequences (5’-TTTCTGTTGGTGCTGATATTGC-forward primer-3’, 5’-ACTTGCCTGTCGCTCTATCTTC-reverse primer-3’) to ensure they were compatible for downstream PCR indexing with the 96 PCR Barcoding Expansion Kit (EXP-PBC096, Oxford Nanopore Technologies, UK).

Both primer sets were applied to all eDNA samples in separate PCRs. Initial PCRs were performed with a final volume of 25 µL, containing 12.5 µL of QIAGEN Multiplex PCR Master Mix (Qiagen, Germany), 0.75 µL of each primer (10 µM), 9.5 µL of ddH2O and 1.5 µL of eDNA on a T100 Thermal Cycler (BioRad, USA). PCR conditions started with an initial denaturation at 95 °C for 15 min, followed by 35 cycles of denaturation at 95 °C for 30 s, annealing at 55 °C for 30 s, and elongation at 72 °C for 30 s, followed by a final elongation at 60 °C for 10 min. Amplification success was confirmed by electrophoresis in 2 % agarose gels stained with GelRed (Biotium, USA). Only successful amplifications were selected for further steps.

Libraries were prepared following the Oxford Nanopore Technologies “PCR barcoding (96) amplicons (SQK-LSK109) (version: PBAC96_9069_v109_revN_14Aug2019)” protocol, with the following specifications: QIAGEN Multiplex PCR Master Mix (Qiagen, Germany) was used for the barcoding PCR; 5 µL of each barcoded PCR product were pooled; fragment size distribution was measured using the 2200 Tapestation System (Agilent Technologies, USA); the barcoded pool was purified using Agencourt AMPure XP beads (Beckman Coulter, USA) using a 0.6 X ratio; pool concentration was measured using Qubit fluorometer with the dsDNA BR Assay Kit (Thermo Fisher Scientific, USA). To facilitate future research the full detailed laboratory protocol is provided (see Data Accessibility).

The final pool was sequenced on a MinION sequencer (Mk1B; Oxford Nanopore Technologies, UK) using an R9.4 flow cell (FLO-MIN106D; Oxford Nanopore Technologies, UK). Starting flow cell pore availability was 908 and the run lasted c. 9 h. It should be noted that samples from another project (using different indexing barcodes, different primers and targeting different genes) were also included in the sequencing run.

The costs and hands-on effort of laboratory sample processing were compared to that which would be expected for an analogous Illumina Miseq protocol. The latter was based on the usual protocols used by the EnvMetaGen group at CIBIO-InBIO (Portugal), which are described by Paupério et al. (2018), Egeter et al. (2018) and Egeter et al. (2019), but using Illumina Nextera XT 96 indexes to allow fair comparison. Sequencing costs were based on the average reads per sample obtained in this study. Costs for DNA extraction were excluded as they are independent of the platforms used. Effort required was estimated for hands-on laboratory time only, conducted by an experienced technician.

### Data processing

Basecalling and demultiplexing were performed using Guppy (v3.4.4; ONT; high accuracy base calling mode). As reads produced by the MinION sequencer are known to have a higher error rate than reads produced by other short read sequencing technologies, two data processing pipelines were considered and tested: one tries to reduce the error of the reads by polishing them after basecalling (henceforth Polished pipeline); the other pipeline skips the polishing, using the raw reads produced after basecalling (henceforth Raw pipeline).

Reads with high error rates could cause issues with taxonomic assignment. One way to mitigate this potential issue would be to restrict the reference database to only those taxa known to be present in the study area and to choose suitable thresholds for assigning taxonomy. This would avoid obtaining matches to species not present in the study area. However, it is not always possible to know *a priori* which species are present. To investigate potential database effects, two effective databases were used separately for taxonomic assignment: The first was the NCBI nt database (accessed April 2020), limited to the taxon Bivalvia (NCBI taxid 6544), (henceforth “bivalves_DB”). The second was the same nt database, limited to the eight species that are potentially present at the study sites (henceforth “8targets_DB”). Additionally, the entire nt database was used, but only for the Polished pipeline, as it would be very time-consuming using the Raw pipeline.

Both pipelines included a step to filter sequences based on length. The expected insert length was obtained by running ecoPCR (Ficetola et al., 2010) on 16S bivalve sequences extracted from the nt database (max error = 3; the sequences used for this step are available in Biostudies, see Data Accessibility). For the VENE primer set, mean insert length was 139bp and although the majority of in silico amplified sequences (92%; 3556/3875) had inserts between 100 and 200bp, insert length ranged from 69bp (family: Veneridae) to 605bp (family: Unionidae). Similarly, for the UNIO primer set, mean insert length was 135bp and although the majority of in silico amplified sequences (95%; 4902/5184) had inserts between 100 and 200bp, insert length ranged from 65bp (family: Veneridae) to 505bp (family: Chamidae). The full ranges were used for length filtering.

The Raw pipeline consisted of two main steps: 1) Primers were trimmed using cutadapt (v2.7; Martin, 2011) accepting a maximum error rate of 20%. We considered that the linked primers could be in either 5’-3’ or 3’-5’ orientation. Reads that did not contain linked primers were discarded; 2) Reads outside the expected amplicon length range for each primer were discarded. The Polished pipeline consisted of four main steps: 1) reads shorter than 40 bases were discarded using cutadapt (v2.7; Martin, 2011); 2) the remaining reads were clustered with isONclust (v0.0.6; Sahlin & Medvedev, 2019) and then polished using racon (v1.3.3; Vaser, Sović, Nagarajan, & Šikić, 2017); 3) the polished reads were then clustered at 99% sequence identity with cd-hit (v4.8.1; Fu, Niu, Zhu, Wu, & Li, 2012); 4) primer trimming and length filtering of the polished reads was performed in an identical manner to the raw pipeline. The Polished pipeline was compiled into a bioinformatics package, *msi* (v0.2.9), which can be easily used in future metabarcoding studies that utilise nanopore sequencing (see Data Accessibility).

For each database, and for each pipeline, the reads (raw and polished) were aligned against the reference database using BLAST (blastn algorithm, v 2.10.0). To ensure that alignments covered at least 98 % of the query sequence, and that multiple hits were returned for each query, we used the following settings: -word_size 11 -perc_identity 50 -qcov_hsp_perc 98 – gapopen 0 -gapextend 2 -reward 1 -penalty −1 -max_target_seqs 50. Following BLAST, taxonomy was assigned to each query using a lowest common approach, similar to (Chain, Brown, MacIsaac, & Cristescu, 2016; Egeter et al., 2018; Kitson et al., 2019): for each query relatively stringent percentage identity thresholds were used – 99 % for species level, 97 % for genus level, 95 % for family level and 93 % for higher-than-family level; for each threshold the lowest common ancestor of remaining alignments was obtained. This software developed for this step was compiled into a separate bioinformatics package, *metabinkit* (v 0.1.3; see Data Accessibility) that is also available for future studies. Finally, to remove any potentially spurious detections, detections with a read count < 50 were removed.

### Statistical analysis

All statistical analysis was performed in R (v3.6.0; R Core Team, 2019). The proportion of reads assigned at each taxonomic level using either the Raw or the Polished pipeline was compared using a two-sample test of proportions (prop.test function in base R). The same test was performed to compare the proportions of reads in each pipeline with top percentage identities in the following categories: 99-100%, 95-97%, 90-95%, 80-90%,70-80% and 50-70%. Heatmaps were prepared using the package ComplexHeatmap (v2.2.0; Gu, Eils, & Schlesner, 2016).

## Results

Most PCRs performed on DNA samples from the Italian sites had evident amplification using the VENE primer set (78 %) whilst none of the Portuguese samples exhibited amplification (see Supporting Information, Table S1 for details of each sample). Similarly, using the UNIO primer set, 73 % of samples from the Italian sites showed amplification, but only one sample collected from Portugal amplified. None of the field negatives or PCR negatives showed any evidence of amplification.

Based on the number of reads obtained for the full sequencing run (c. 5.3 million), the estimated flow cell run time for libraries that were part of this project was 1.6 h (total 9 h). The estimated time for basecalling and demultiplexing was 1.5 h (total 8 h), yielding 652,258 and 307,062 reads for VENE (n PCR products = 14) and UNIO (n PCR products = 13), respectively. The number of reads at each bioinformatic step are provided in Supporting Information Table S2. Processing time was much faster using the Polished pipeline than using the Raw pipeline (Table 2). This was mainly due to the BLAST step, which for the Raw pipeline essentially involved mapping all reads against the reference database, while for the Polished pipeline involved aligning only the operational taxonomic units (OTUs).

**Table 2.**
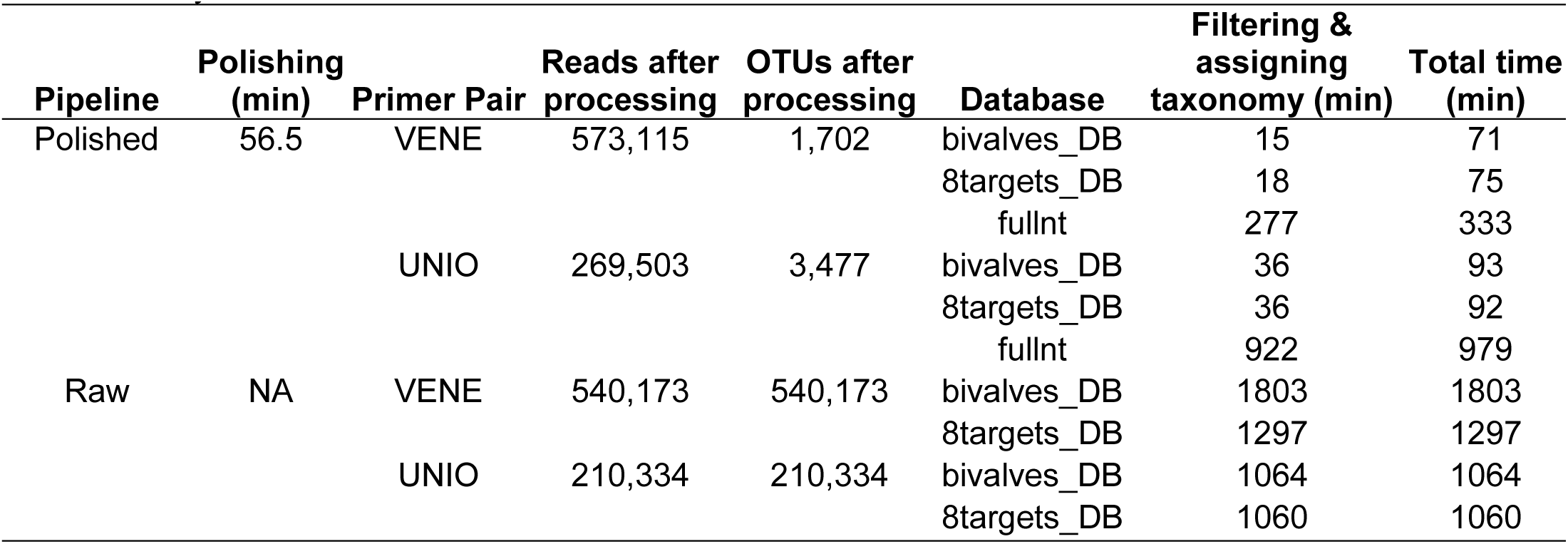
Number of reads and OTUs after processing, and CPU processing time required for each pipeline and database. Note that primer trimming is included in the Polishing step for the Polished pipeline, but is included in the Filtering & assigning taxonomy step for the Raw pipeline. All analyses were conducted on the same machine.

Regardless of the database used, the percentage identities of the top BLAST hits were significantly higher using the Polished pipeline (Figure 3; Supporting Information Table S3; p<0.001 for all categories apart from the 70-80% category using the 8targets_DB, where p=0.006). The largest category of top BLAST hits using the Raw pipeline was 90 – 95 % identity (45 % of reads using the 8targets_DB and 42 % of reads using the bivalves_DB), whilst using the Polished pipeline the largest category was 99-100 % identity (62 % of reads using the 8targets_DB and 60 % of reads using the bivalves_DB).

**Figure 3.**
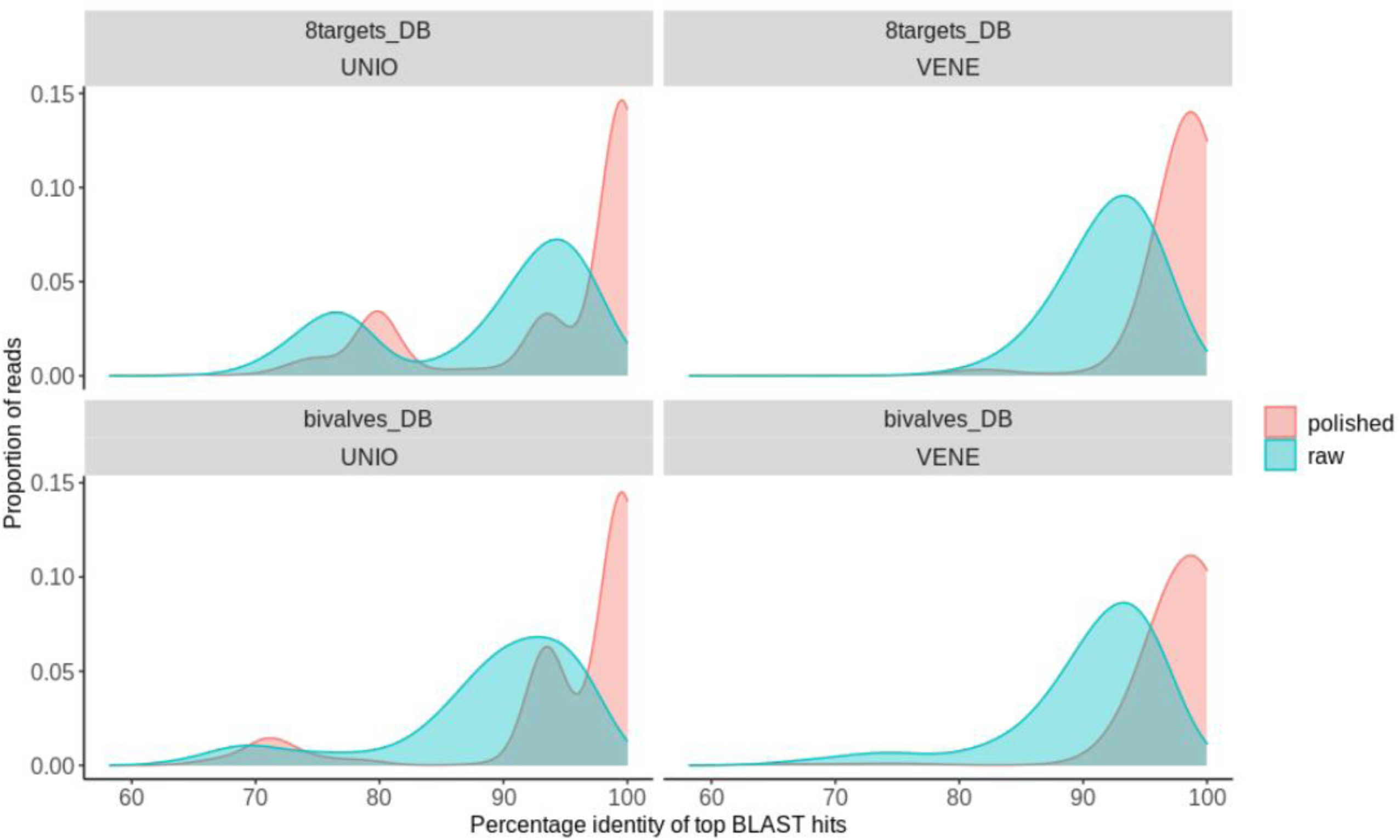
The relationship between proportion of reads and top BLAST hit percentage identities for each pipeline and database combination. See Supporting Information Table S3 for exact proportions in each category, as well as associated chi-square and p-values.

Using identical BLAST and taxonomic assignment procedures, the proportion of reads being assigned at each taxonomic level was always significantly higher (p<0.001 in all cases) using the Polished pipeline (Figure 4; Supporting Information Table S4). Using the Polished pipeline and the 8targets_DB, 67 % of reads were assigned at species level, in contrast to 2 % using the Raw pipeline. Using the bivalves_DB 37 % of reads were assigned at species level using the Polished pipeline, in contrast to 1 % using the Raw pipeline (Supporting Information Table S4).

**Figure 4.**
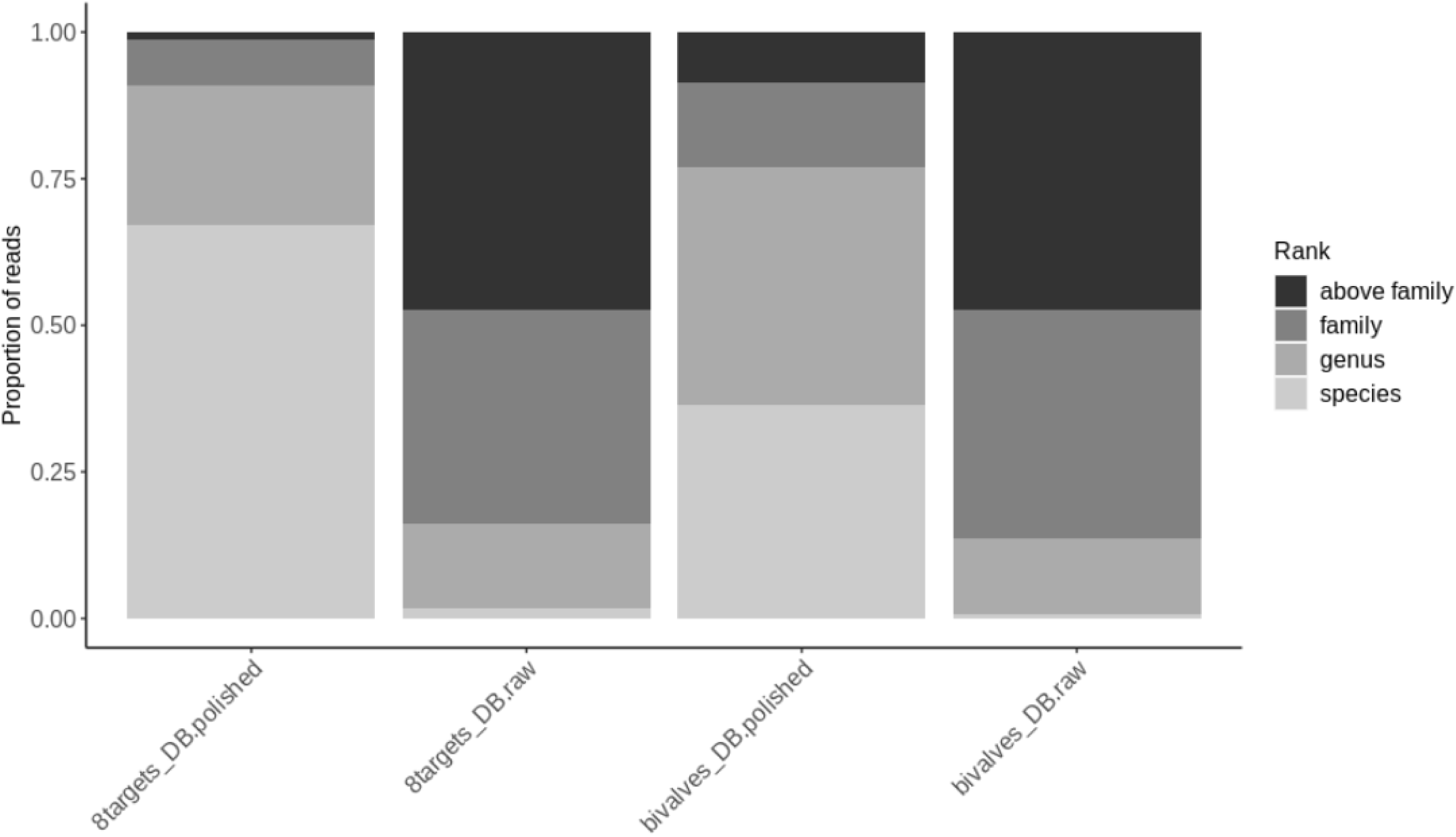
The proportion of reads assigned to each taxonomic level for each pipeline and database combination. See Supporting Information Table S4 for exact proportions in each group, as well as associated chi-square and p-values.

There were no detections in the Raw pipeline that were not also in the Polished pipeline. All species-level detections in the Polished pipeline had higher read counts than those in the Raw pipeline. The Polished pipeline resulted in detections that were not observed in the Raw pipeline. There were no additional bivalve species or detections using the full nt or the bivalves_DB rather than the 8targets_DB. Notably, although using the full nt database took considerably longer (Table 2), additional non-bivalve taxa were detected (Supporting Information Figure S1), including bacteria, mayfly (*Cloeon dipterum*), fish (*Salaria fluviatilis, Lepomis gibbosus, Micropterus salmoides*), and birds (*Cygnus olor, Fulica atra, Podiceps cristatus*), all of which are taxa known to occur at the sites sampled. Taking these points into consideration, the following results focus only on the data obtained using the Polished pipeline and the 8targets_DB. The combined final taxa table and a side-by-side heat map produced by all pipeline-database combinations are provided in the Supporting Information (Table S5, Figure S1).

In total, five species of bivalve were detected, belonging to four genera from three families (Figure 5). No taxa were detected at Portuguese sites. All three expected invasive species and two of the four expected native species were detected at Italian sites. The invasive *D. polymorpha* was detected at all Italian sites and at none of the Portuguese sites. At Lake Maggiore and Lake Varese, in Italy, two further invasives, *C. fluminea* and *S. woodiana*, and one native species, *U. elongatulus*, were detected. One further native species, *A. exulcerata*, was detected only at Lake Varese. As none of the species detected in Italy are known to be present at the sites in Portugal, the lack of detections from Portuguese sites indicates that there was no contamination between samples, either physically or due to demultiplexing or taxonomic assignment errors. Almost all absences of species detections can be explained either by the known species occurrences or by the fact that they are known to occur at low densities at the respective site (Figure 5). The only possible exception is an absence of detection of *C. fluminea* at Lake Lugano, which is known to occur at that site and can be at reasonably high densities (c. 2 individuals per m^2^).

**Figure 5.**
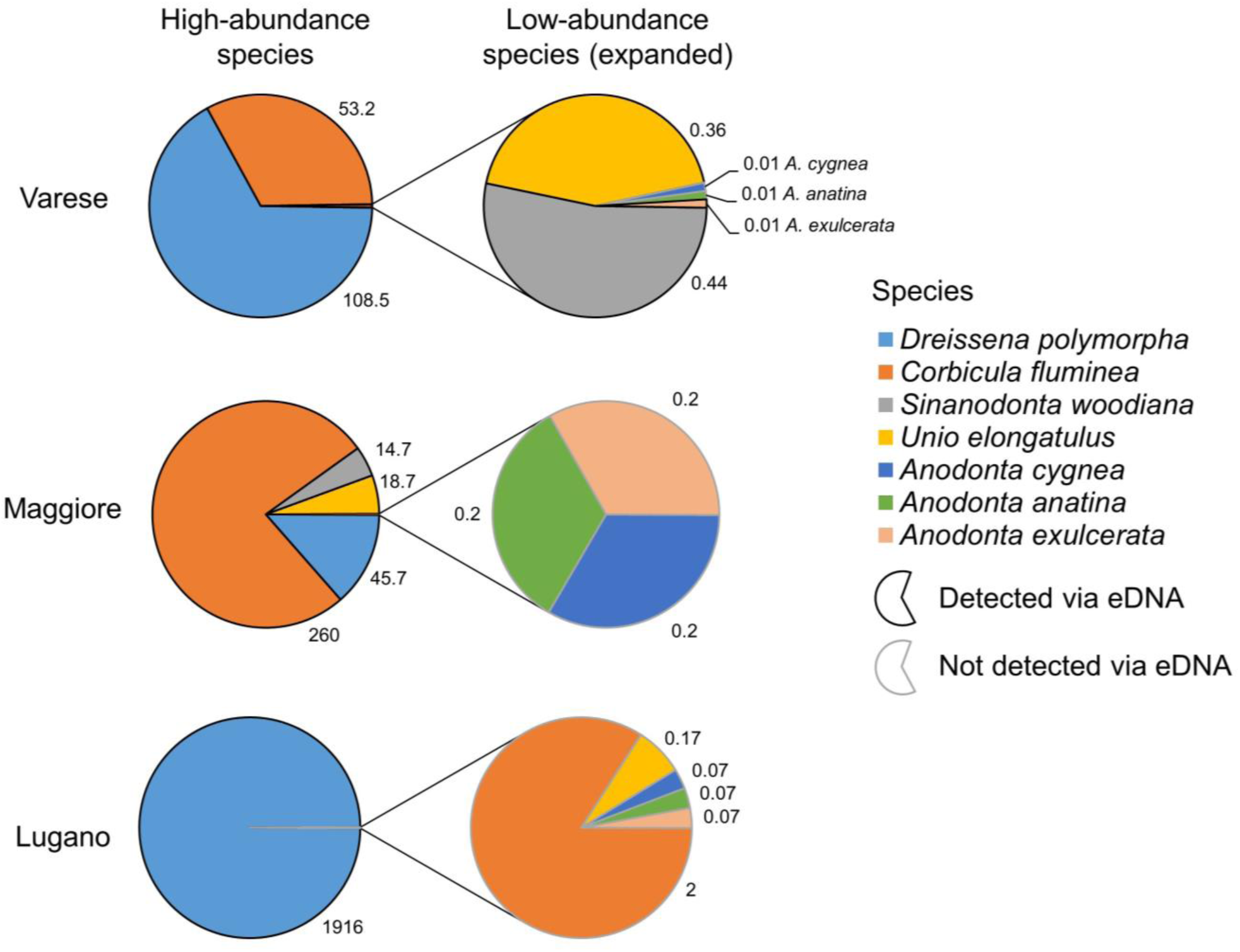
The maximum abundance (individuals/m^2^) of each bivalve species at each Italian site observed during surveys in 2019 (Supporting Information Table S6). Black-highlighted segments indicate the species detected in the present study, from at least one sample. Grey-highlighted segments indicate the species not detected in the present study. See Supporting Information Figure S1 for sample-level details.

The laboratory costs and hands-on effort of using the MinION protocol were very similar to that which would be expected for an analogous Illumina Miseq protocol (Table 3).

**Table 3.**
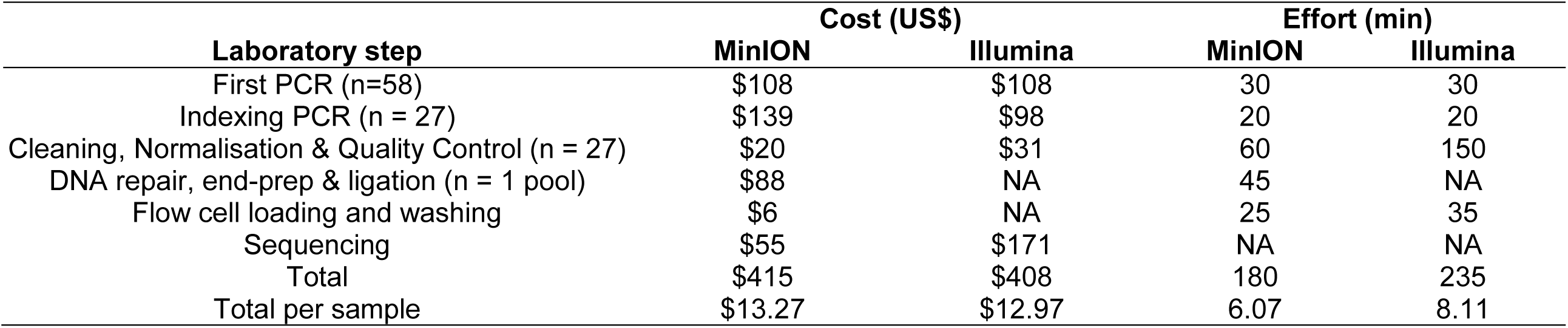
Estimated laboratory costs and effort required for this study 1) using the MinION protocol described herein and 2) had an analogous Illumina Miseq protocol been used instead.

| Laboratory step | Cost (US\$) | | Effort (min) | |
| --- | --- | --- | --- | --- |
|  | MinION | Illumina | MinION | Illumina |
| First PCR (n=58) | \$108 | \$108 | 30 | 30 |
| Indexing PCR (n = 27) | \$139 | \$98 | 20 | 20 |
| Cleaning, Normalisation & Quality Control (n = 27) | \$20 | \$31 | 60 | 150 |
| DNA repair, end-prep & ligation (n = 1 pool) | \$88 | NA | 45 | NA |
| Flow cell loading and washing | \$6 | NA | 25 | 35 |
| Sequencing | \$55 | \$171 | NA | NA |
| Total | \$415 | \$408 | 180 | 235 |
| Total per sample | \$13.27 | \$12.97 | 6.07 | 8.11 |

## Discussion

Our study clearly showed the value of our approach to reliably detect multiple invasive species using nanopore sequencing technology. We suggest that using nanopore sequencing technology combined with appropriate bioinformatic pipelines can provide a reliable, inexpensive and rapid method for researchers and environmental managers to detect invasive species, and could be widely adopted for applications requiring the targeted detection of single or multiple species (e.g., endangered species, disease vectors, and economically valuable species).

### Reliability and resolution

Although MinION reads have a much higher error rate than those produced by Illumina platforms, it was possible to overcome this using the Polished pipeline. This is supported by the following evidence: 1) 62 % of the polished reads were 99-100 % identical to sequences in the reference databases; 2) there were no taxa detected other than those suspected to be present within the study sites (i.e. no false positives); 3) using relatively stringent taxonomic assignment thresholds, 99 % of reads could be assigned to family level, 91 % to genus level and 67 % to species level (using the 8targets_DB).

One notable difference between the bivalve_DB and the 8targets_DB analysis was that *C. fluminea* could not be assigned at species level using the bivalve_DB due to very similar matches in the database to other *Corbicula* species (primarily *C. colorata* and *C. leana*). The authors of the primer sets used in the current study noted that taxonomy is still unclear for this genus, that GenBank specimen determination may be hazardous, and that the *Corbicula* species complex cannot currently be distinguished with this method (Prié et al., 2020). Additional reference 16S sequences are required, from carefully identified specimens, or other genes need to be targeted to enable species-level detection of this genus in future eDNA studies.

### Detection sensitivity

The lack of detections of each species at each site could be explained either by its known absence from the site or because the species is known to occur only at low densities. The most likely reason for not detecting species that occur at low densities is that eDNA of these species is likely to be present in very low concentrations, particularly in relation to species that occur at high densities. For example, *C. fluminea* was not detected at Lake Lugano although it occurs here at up to 2 individuals/m^2^, but at this site *D. polymorpha* occurs at extremely high densities (815 – 1916 individuals/m^2^). Thus it is likely that PCR competition and/or insufficient sequencing depth likely prevented the detection of this proportionally less abundant taxon (see Elbrecht, Peinert, & Leese, 2017). We also noted that there was considerable variation in the number of reads obtained for each PCR (range: VENE 10k – 84k; UNIO 2k – 70k), which is likely to exacerbate sequencing depth issues for samples with lower read numbers. While we did pool barcoded PCRs in equal volumes, we did not pool with equal molarity. We suggest this is carried out in future MinION studies.

Given that the error rates in MinION reads can be overcome, the factors that will most influence the sensitivity of an eDNA approach for the detection of aquatic invasive species are likely to be external to the utilisation of nanopore sequencing technology, i.e. using optimal field sampling methods, extraction protocols and PCR conditions, along with choosing suitable primers, sequencing to a sufficient depth and conducting multiple PCR replicates. All of these factors have been shown to increase the detection probability of low abundance taxa (Nichols et al., 2018; Shaw et al., 2016; Zaheer et al., 2018), and are particularly important in cases where sympatric species occur in high densities. As noted by Hatzenbuhler et al. (2017), practical early detection strategies need to balance search effort with an acceptable amount of non-detection risk. The effort required to conduct MinION-based detection of aquatic invasive species is far lower than using traditional morphological taxonomy, is on par with other eDNA approaches, and has the important benefit of reduced turnaround times.

### Efficiency

In the current study, it was possible to extract, amplify, prepare, and sequence DNA from 27 successful PCRs with a turnaround of c. 3.5 days. Truelove et al. (2019) also reported short MinION turnaround times (c. 48 h) for detecting sharks from water samples while aboard a ship, although they did not attempt barcoding of multiple PCR products, so their turnaround times are for each sample independently. One of the major benefits of using the MinION for metabarcoding over the Illumina platform is the ability to terminate runs before a flow cell is exhausted, allowing it to be used for multiple runs. This is especially valuable when sample sizes are low, as a pool of just a few samples can be prepared, loaded and sequenced independently, without having to wait for additional samples to become available to fill a run. In contrast, for Illumina-based approaches it is common that smaller projects are shelved until there are enough libraries prepared from other projects to justify using a full Miseq / Hiseq flow cell. Furthermore, as the MinION itself is relatively inexpensive (US$1,000 at the time of writing, including two flow cells), it avoids the need to contract external sequencing services, which often have turnaround times of 2-4 weeks (and in some cases can take much longer).

The costs of the indexing PCR step using the PCR Barcoding Expansion Pack 1-96 (Oxford Nanopore Technologies; EXP-PBC096) at the time of writing equates to US$1.73 / PCR. This cost could be substantially reduced by using custom-made barcodes. For example, Srivathsan et al. (2019) estimated their cost of producing DNA species barcodes as US$0.35 per barcode, by incorporating 13bp tags in their primers and multiplexing c. 3,500 samples per flow cell. Such approaches also increase the scalability of using MinION for studies with larger sample sizes. Based on other MinION runs performed in our laboratory (EnvMetaGen group, CIBIO-InBIO, Portugal) that all comprised relatively short metabarcoding amplicons, we obtain an average of 0.4 million reads/h on a flow cell that can be used for up to 48 h, equating to c. 19 million reads per flow cell, making the cost of the sequencing step cheaper using MinION than it would be using the Illumina Miseq. Of course, this will vary according to the number of available and actively sequencing pores on the flow cell throughout the run time.

### Applications

Overall, this approach proved to successfully detect most of the invasive bivalve species in the study sites, without the requirement of morphological taxonomic identification or intensive survey methodologies. For instance, the detection of *D. polymorpha* in southern Portugal (Catita et al., 2020) required the deployment of monitoring ropes in several water bodies. This approach is logistically challenging and has a high turnaround time between sample collection (i.e. rope deployment and consequently colonization) and species detection. Approaches utilising eDNA and nanopore technology have the potential to considerably reduce such field survey requirements and turnaround times.

Nanopore technology has the ability to sequence much longer reads than Illumina technology, and our sequencing run did result in reads over 600 bp, after primer trimming. In the case of this data set, however, these reads had no significant alignments in the bivalve databases and upon further inspection mapped most closely to bacterial sequences. We did not specifically target longer amplicons in this study, as detection of faunal eDNA is negatively affected by amplicon length (Bylemans, Furlan, Gleeson, Hardy, & Duncan, 2018; Rees et al., 2014). However, some eDNA studies have successfully sequenced eDNA fragments longer than the usual 50-250 bp fragments (e.g. Deiner et al., 2017; Ma et al., 2016) and it is likely that future studies could avail of nanopore technology to sequence such amplicons.

Another potential benefit of the MinION sequencer is the ability to sequence DNA using a portable laboratory (Maestri et al., 2019; Pomerantz et al., 2018; Truelove et al., 2019). Although we did not attempt this in the current study, the laboratory protocols and bioinformatic software that we provide could be adapted to such conditions, as they do not necessarily rely on bulky equipment or an active internet connection. This would allow results to be obtained quickly on site, or in field accommodation, and has the added benefit of avoiding potential laboratory-derived contamination.

Previous studies have demonstrated the application of nanopore technology for generating DNA barcodes and identifying species from tissue-extracted DNA (Ho et al., 2020; Krehenwinkel et al., 2019; Seah et al., 2020), assessing microbial diversity from drinking water sources (Acharya et al., 2019) and detecting shark species from marine samples (Truelove et al., 2019). The fact that we also identified bacteria, vertebrate and insect taxa (Supporting Information Figure S1), provides further evidence that the nanopore sequencing could be adapted for many of the already existing eDNA applications (e.g. biodiversity assessment, ecosystem health assessment, diet studies).

### Conclusion

Invasive species can be difficult to eradicate once established, therefore continuous monitoring in un-invaded systems is crucial for the quick detection and successful suppression of these species. In general, early detection of aquatic invasive species increases the probability that control and eradication efforts will be successful (Anderson, 2005; Goldberg, Sepulveda, Ray, Baumgardt, & Waits, 2013). We present a reliable, cost-effective approach for detecting aquatic invasive species that reduces turnaround time in data acquisition and can be easily adapted for other metabarcoding studies. To facilitate future research we provide a detailed library preparation protocol as well as an open source software package that wraps the computational analysis workflow.

## Acknowledgments

This project has received funding from the European Union’s Horizon 2020 Research and Innovation programme under grant agreement No 668981 (ERA Chair in Environmental metagenomics, EnvMetaGen), EDP-Biodiversity Chair (EDP/FCT), and by the project PORBIOTA-Portuguese E-Infrastructure for Information and Research on Biodiversity (POCI-01-0145-FEDER-022127), supported by Operational Thematic Program for Competitiveness and Internationalization (POCI), under the PORTUGAL 2020 Partnership Agreement, through the European Regional Development Fund (FEDER). JV is supported by FCT PhD grant SFRH/BD/133159/2017. We would like to thank everyone in the EnvMetaGen team for their support and advice during the development of this study.

## Data Accessibility

All data and analysis scripts underlying this study are accessible via BioStudies Accession Number S-BSST391 (https://www.ebi.ac.uk/biostudies/studies/S-BSST391). The raw sequencing data has been deposited at the European Nucleotide Archive (ENA) under the accession number PRJEB38199 (https://www.ebi.ac.uk/ena/browser/view/PRJEB38199). The detailed laboratory protocol developed as part of this study, *Metabarcoding using MinION: PCR, Multiplexing and Library Preparation*, is available on protocols.io (https://dx.doi.org/10.17504/protocols.io.bfqqjmvw). The software packages developed as part of this study are publicly available: *msi* (https://doi.org/10.5281/zenodo.3872794) and *metabinkit* (https://doi.org/10.5281/zenodo.3873646). A data package has also been provided which can be used to rerun the polishing and taxonomic assignment analyses performed in this study (https://github.com/envmetagen/mussels_package).

## Author Contributions

PB, NAF and BE conceived the study. JV, MLL and NR collected field samples. BE, JV, CC and JP contributed to the development of the laboratory protocol. CC and JP conducted the laboratory sample processing. NAF and BE wrote the bioinformatics software and processed the data. BE conducted the statistical analyses and led the manuscript preparation. All authors contributed to the preparation of the manuscript.

